# Landowner Functional Types to Characterize Response to Forest Insects

**DOI:** 10.1101/596189

**Authors:** Jonathan R. Holt, Mark E. Borsuk, Brett J. Butler, David B. Kittredge, Danelle Laflower, Meghan Graham MacLean, Marla Markowski-Lindsay, David Orwig, Jonathan R. Thompson

## Abstract

Forest insects and pathogens (FIPs) play an important role in the complex interactions between woodland owners and the ecosystems they manage. Understanding the specifics of woodland owner decision-making with regard to trees impacted by FIPs can facilitate projections of future forest conditions and insect spread. Our first objective is to: (i) characterize agent functional types (AFTs) of New England family forest owners (FFOs) using a set of contingent behavior questions contained in a mail survey of FFOs concerning response to FIPs. We establish AFTs as a form of dimension reduction, effectively assigning individual FFOs to particular decision-making classes, each with distinct probabilities of behavior with respect to the harvest of impacted trees. Our second objective is to: (ii) model AFT membership to predict the distribution of AFTs across the landscape. Predictors are chosen from a set of survey, geographic, and demographic features. Accomplishing (i) establishes three functional classes of landowners: ‘cutters’ (46% of respondents; highly likely to cut timber impacted by FIPs), ‘responsive’ cutters (42% of respondents; sensitive to pest severity), and ‘non-cutters’ (12% of respondents; highly unlikely to cut). Fulfilling (ii) provides a high-resolution probability surface of AFTs across the landscape, delivering key input for simulation models of forest and land cover change in New England. Predictors in our best model of AFT membership include parcel size (hectares of forest), region, and town-level forested fraction.

## 1. Introduction

Invasive forest insects and pathogens (FIPs) are a prominent cause of forest disturbance in North America (Thorn et al., 2018). The U.S. Forest Service’s National Insect and Disease Forest Risk Assessment (Krist Jr et al., 2014) suggests that 334 million ha, or 63% of the nation’s forestland, are at risk for host basal area mortality, and 24.8 million ha are predicted to experience more than 20% loss of host basal area through 2027. In fact, FIPs are the only forest disturbance agent that has proved capable of nearly eliminating entire tree species, or in some cases entire genera, within a matter of decades (Lovett et al., 2016).

While the direct effects of FIPs on tree mortality are relatively well understood, the indirect impacts of FIPs via “pre-emptive” or “salvage” harvesting are not as well studied (Foster & Orwig, 2006). Harvesting is currently a larger cause of mature tree mortality in northeastern forests than all others combined (Canham, 2013); the frequency and intensity of harvests varies widely depending on both biophysical and social factors (Thompson, Canham, Morreale, Kittredge, & Butler, 2017). Past FIP outbreaks in northeastern forests have been accompanied by accelerated harvesting, and there are distinct ecological legacies of the interactions between these two types of biotic disturbance (Thorn et al., 2018). For example, following reports that hemlock woolly adelgid (*Adelges tsugae*) had reached Connecticut in the 1980s and 1990s, many landowners harvested hemlock trees, despite their low commercial value (Orwig, Foster, & Mausel, 2002). Since the disturbance from harvesting in response to FIPs can be more intense than that of the FIP alone (Foster & Orwig, 2006), there is a need to better understand when and why landowners harvest in response to FIPs.

FIPs present forest owners with the risk of diminished economic returns and/or woodland aesthetics (Li et al., 2014). How a landowner responds to this risk can be expected to depend on his or her objectives in owning their forest (Nordlund & Westin, 2011), as well as socioeconomic and demographic factors. The anticipated severity of the outbreak, the location of the woodland, and FIP awareness may also be factors that influence the landowner’s response (or lack thereof) to FIPs (Boyd, Gilligan, & Godfray, 2013; Nielsen-Pincus, Ribe, & Johnson, 2015). To better understand these influences, we surveyed private, non-industrial, family forest owners (FFOs) in New England (northeastern United States) (Markowski-Lindsay et al., 2019). New England is an ideal study system because the region contains many private landowners and one of the highest diversities of FIPs in North America (Liebhold et al., 2013). In New England, an estimated 41% of all forestland is controlled by FFOs (B. J. Butler et al., 2016). Thus, the response of FFOs to FIPs may intensify, broaden, and potentially synchronize the ecosystem impacts of FIPs on the landscape.

Our survey contained contingent behavior questions (CBQ) to assess whether an FFO would harvest in response to different FIP scenarios. We used the responses to the CBQ to cluster individuals into agent function types (AFTs). The idea is to group together similar “types” of people based on common behavior. This form of dimension reduction leads to a functional typology which, while not describing the individual, represents archetypal patterns of behavior that tend to repeat themselves within the community (Ficko, Ní Dhubháin, Karppinen, & Westin, 2019). AFTs have proven to be useful for modeling human decision-making in a variety of applications, especially in the agricultural sector (Guillem, Barnes, Rounsevell, & Renwick, 2012; Karali, Brunner, Doherty, Hersperger, & Rounsevell, 2013) and in the context of large-scale socio-ecological systems (Arneth, Brown, & Rounsevell, 2014; Rounsevell, Robinson, & Murray-Rust, 2012). They have been less frequently used to represent private woodland owners (except e.g. Blanco et al., 2015). Further, the majority of landowner typologies have been based on objectives for ownership (Kelly, Gold, & Di Tommaso, 2017; Khanal et al., 2017; Nielsen-Pincus et al., 2015) or more nuanced criteria such as attitudes towards climate change (Khanal et al., 2016), approaches to fire management (Charnley, Kelly, & Wendel, 2017), and thoughts on pollution (Perry-Hill & Prokopy, 2014). These are all proxies for behavior rather than explicitly functional behaviors.

In the present study, we form AFTs based on landowner responses to our survey CBQ. In this way, the contingent *behaviors* revealed by the survey responses generate a *functional* typology. We then develop a model to predict AFTs for FFOs who did not participate in the survey, giving us a means for scaling the survey data to the entire region and yielding probability distributions for the various AFTs across the landscape. Ultimately, this AFT framework allows us to estimate the probability that an FFO will cut their trees in response to a particular FIP scenario. Understanding such harvest probabilities across the region provides insight into the condition in which FFP response may intensify the impacts of FIPs on our forests.

## 2. Data & Methodology

### 2.1 Overview

To comprehensively understand landowner response to FIPs, we used data from our woodland owner survey to: (i) fit a binary logistic regression model to the contingent behavior results thereby relating harvest probability of respondents to various FIP conditions; (ii) group FFOs into AFTs based on the observed patterns of contingent behavior; (iii) compare and contrast AFTs with respect to survey-reported age, education, income, and other household-level demographics; (iv) characterize AFTs with respect to reported management history and objectives for ownership; (v) identify socio-demographic differences across AFTs at the town-level using the American Community Survey (Manson, Schroeder, Van Riper, & Ruggles, 2017); and (vi) develop a multinomial logistic regression model that predicts AFT membership as a function of geographic features, allowing us to map AFT probabilities across the landscape and thereby expose spatial patterns of landowner responses to FIPs.

### 2.2 Landowner Survey

We designed and administered the New England Woodland Owner Survey (NEWOS), which was mailed to a random sample of FFOs owning ≥ 4ha of land in the Connecticut River Watershed (Markowski-Lindsay et al., 2019). The heavily wooded Connecticut River Watershed (Figure 1) lies in the heart of New England, straddling Vermont and New Hampshire to the north and then stretching south through central Massachusetts and Connecticut. Some of the most damaging FIP species in the Connecticut River Watershed include hemlock woolly adelgid (*Adelges tsugae*), emerald ash borer (*Agrilus planipennis*) and European gypsy moth (*Lymantria dispar*). The scenarios presented in the CBQ referenced a generic FIP, but the range of characteristics was chosen based on the three insects mentioned above.

**Figure 1:**
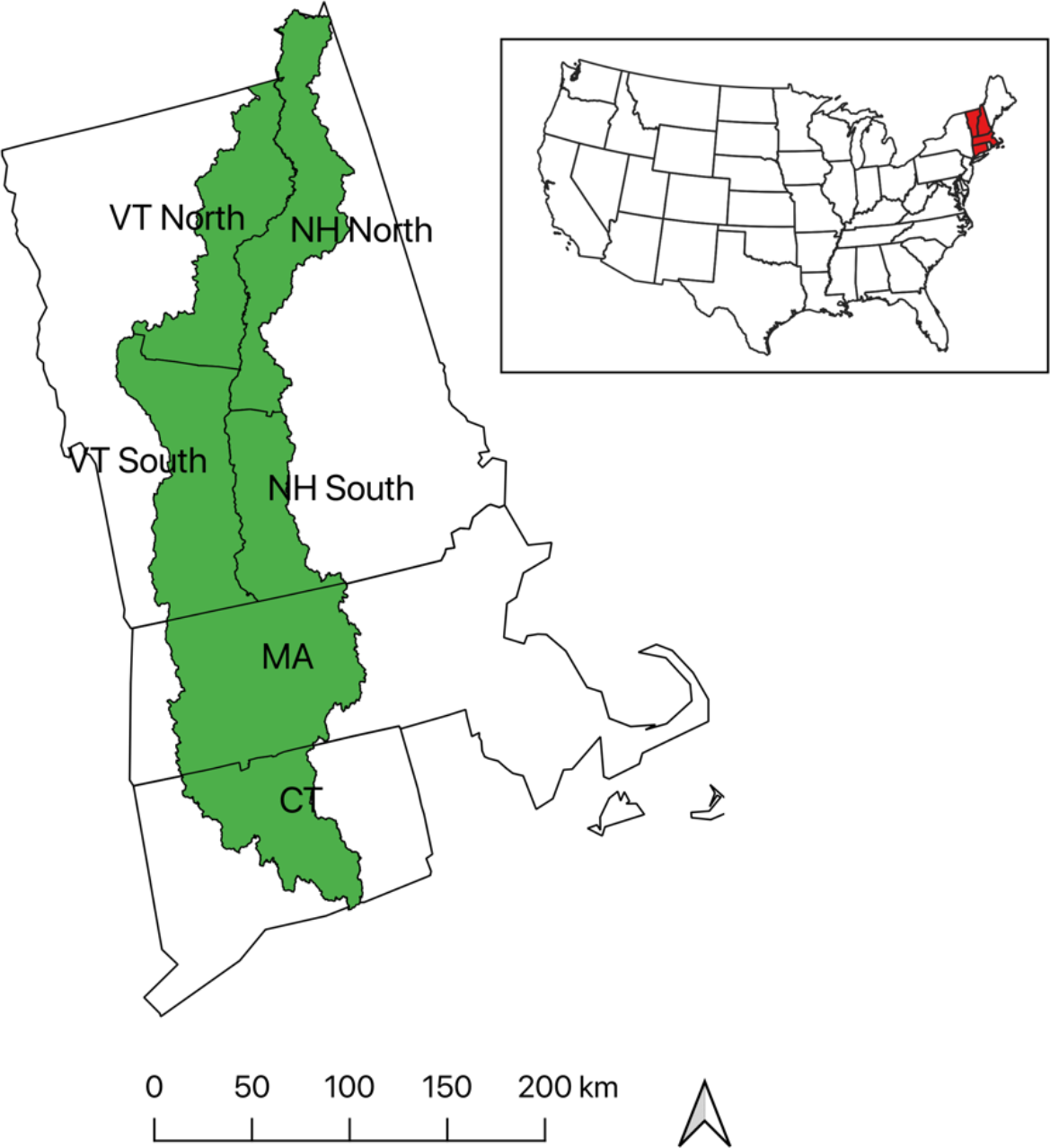
Study area. The Connecticut River Watershed (green) runs through New England in the northeastern United States. The survey was stratified by the six regions shown.

The survey sample was stratified by six regions (New Hampshire north and south, Vermont north and south, Massachusetts, and Connecticut) and also stratified by parcel size (4-19 ha and ≥ 20 ha) to ensure sufficient representation of larger parcels across the region. Landowner demographics, objectives for ownership, familiarity with forest FIPs, and CBQs were among the subsections of the NEWOS. In 2017, 2,000 mail surveys were sent to approximately 333 FFOs per region (roughly 167 per strata). The overall participation rate was 37%, or 688 usable surveys. We detected no nonresponse biases based on telephone follow-up calls or early/late respondent comparisons. See Markowski-Lindsay *et al.*, 2019 for a more in-depth discussion of the survey creation and landowner responses.

The contingent behavior questions in the survey were designed to reveal FFO intended behavior in response to different FIP severity metrics. We presented respondents with a series of scenarios over the range of FIP severity metrics: 1) percent of trees destroyed by the FIP ("mortality percent"); 2) timber value loss due to the FIP ("value loss"); 3) time from now until FIP arrives ("time to arrival"); and 4) time from FIP arrival until tree damage is complete ("time to mortality"). Survey respondents received one of six unique versions of the survey and were presented with five scenarios involving combinations of the four FIP severity metrics (Table 1).

**Table 1:**
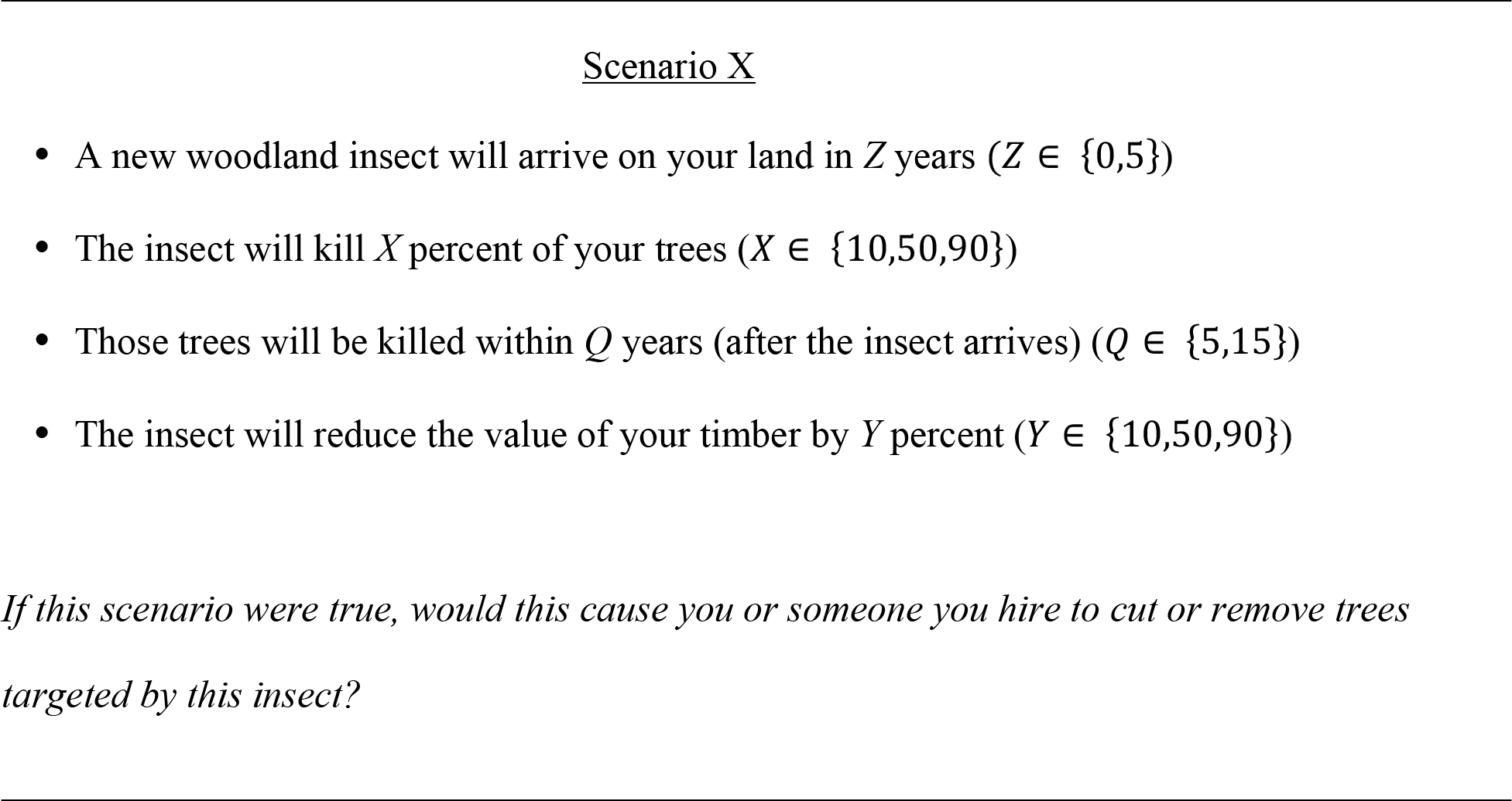
Contingent behavior questions. Responses to each of the five scenarios are binary: yes or no (cut or don’t cut) followed by a measure of respondents’ certainty (five-level Likert scale).

### 2.3 Logistic Regression Model of Contingent Behavior

We fit a binary logistic regression to the responses of FFOs whose answers to the CBQs were sensitive to the stated levels of FIP severity metrics. The resulting model allows us to evaluate the relative importance of each of the severity metrics in determining the likelihood of harvest.

We used a Bayesian model framework, in which *X*, *Y*, *Z*, and *Q* represent the four FIP severity metrics, *i* is the scenario, *j* is the individual, and *y* is the response. Random effects were permitted for the intercept to account for differences across individuals. Priors on the model coefficients *β* were non-informative.

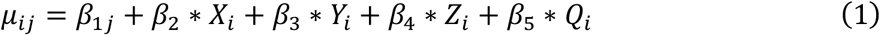

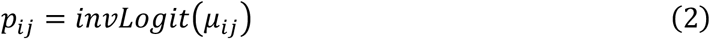

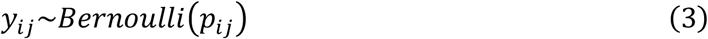

The model was implemented using JAGS via the R2jags package (Su & Yajima, 2012). The model converged after 500 iterations and was run for a total of 1000 iterations. We included survey sampling weights in the regression to generalize to the population; these weights are derived by Markowski-Lindsay et al., 2019.

### 2.4 Characterizing AFTs

Survey respondents were grouped into AFTs based on natural clustering in the responses to the CBQs (details in the Results section). Other survey items were then used to characterize these groups, and the ANOVA posthoc Tukey test was used to identify significant differences. Principal Components Analysis (PCA) was used to downscale two sets of data: 1) landowner objectives from the NEWOS (Supplemental Information A); and 2) socio-demographic features from the ACS dataset. Statistical analyses were conducted in R (R Team, 2017).

### 2.5 Spatial and Demographic Data

Land cover data was obtained from the National Land Cover Database (NLCD) 2011 (Homer et al., 2015). The NLCD land cover classes were consolidated into: “forest” (deciduous forest + evergreen forest + mixed forest + shrub scrub + woody wetlands); “agriculture” (crops + pasture/hay); “low density development” (low intensity development + medium intensity development + open space); “high density development” (high intensity development). Town-level percentages of land cover classes were extracted using town shapefiles and then joined to the NEWOS data.

We obtained town-level socio-demographic data from the 2011-2015 American Community Survey (ACS) (Manson et al., 2017) and joined it to the NEWOS data. Parcel shapefiles were obtained from each state, and occasionally town, individually. Massachusetts was the only state in the region offering a complete parcel dataset. Connecticut, New Hampshire, and Vermont had some missing town parcel data that were filled using towns with similar attributes (size and developed area) from within the study area.

### 2.6 Predicting AFTs Across the Landscape

To predict the spatial distribution of AFTs across the landscape, we developed a multinomial logistic regression model from available spatial data. The full feature set included parcel-scale geographic features (e.g., total area, forested fraction), town-scale geographic features (e.g., land cover), and town-scale socio-demographic features (e.g., median household income). Model selection proceeded in steps: First, we sought to uncover non-linear spatial patterns. To that end, we constructed a classification and regression tree (CART) (Breiman, 1984) with AFT as the response and all spatial parcel data (e.g., total area, latitude/longitude, survey strata) as predictors. Detected patterns were compiled into a new spatial categorical variable (see Appendix B for more information). We then constructed a multinomial logistic regression model with the full feature set, including an interaction term with the spatial categorical variable Finally, we used a stepwise AIC procedure (forwards and backwards) to select the best model. Modeling was conducted in an R environment using the nnet and rpart packages (Ripley & Venables, 2011; Therneau, Atkinson, & Ripley, 2010). The multinomial logistic regression model was then used to predict the probability of each of the AFTs for ~90,000 FFO land parcels throughout the Connecticut River Watershed.

## 3. Results

### 3.1 Logistic Regression Model of Contingent Behavior

Responses to the contingent behavior questions in the survey revealed a natural clustering among respondents. Twelve percent of respondents answered "No" (they would not cut) to all FIP scenarios provided, while 46% of respondents selected "Yes" (they would cut) for all provided scenarios. The remaining 42% of respondents revealed sensitivity in their contingent cutting behavior to variation in the four FIP severity measures. We could thus immediately identify two AFTs: “cutters” – those FFOs who will apparently always harvest in response to FIPs – and “non-cutters” – those who will apparently never harvest in response to FIPs. The remainder of respondents, who we call “responsive cutters”, are evaluated for further subcategorization based on the results of the logistic regression analysis.

Our logistic regression model for the 285 responsive cutters reveals that “mortality percent” (percent of trees killed by a FIP) and “time to mortality” (time from FIP arrival until damage completion) are important predictors of harvest response (Figure 2). The 95% credible interval for the coefficient on mortality percent is entirely positive, indicating that landowners are more likely to harvest trees as the severity of infestation increases. The 95% credible interval for the coefficient on time to mortality is entirely negative, indicating that FFOs are more inclined to harvest when the threat is more imminent. The coefficient on “time to arrival” (time from now until FIP arrives) is largely negative, indicating that landowners are less likely to harvest the more distant the arrival time, but the 95% credible interval includes zero. The coefficient on “value loss” (timber value loss due to FIP) is also primarily negative, indicating a lower likelihood of harvest the greater the value of timber at state, but also has a 95% credible interval that includes zero.

**Figure 2:**
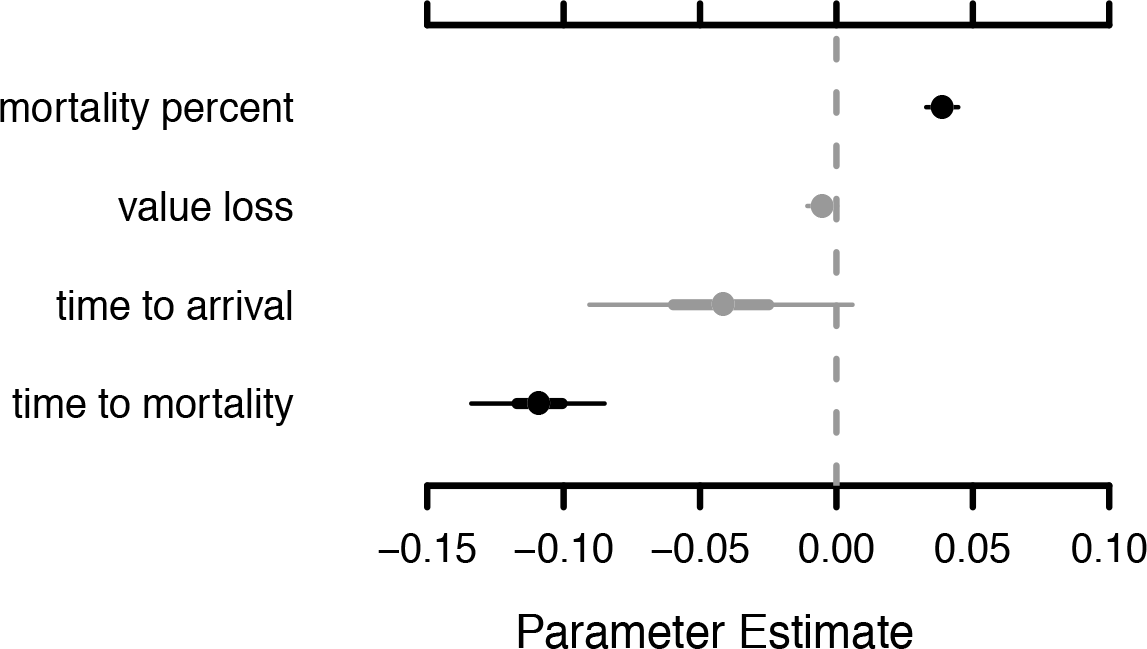
Parameter estimates for a binary logistic regression model fitted to data from the 285 “responsive cutters”. Thicker and thinner lines represent 68 and 95% credible intervals, respectively, with grey lines indicating 95% credible intervals that contain zero.

We attempted to identify additional AFTs from among the group of responsive cutters by performing cluster analysis on the intercepts estimated for each respondent. We also fit logistic regression models in which the coefficients on FIP intensity were allowed to vary by individual and attempted to find natural groupings in these values. While we found that there was some potential variation among the responsive cutters in their baseline tendency to cut (i.e., intercept) and in their sensitivity to intensity measures (i.e., coefficients), clusters that might suggest the need for functional types were not apparent. We therefore proceeded with a total of three AFTs for further characterization.

### 3.2 Characterizing AFTs

Cutters, the most numerous of the three identified AFTs, own on average more area of woodland (55.1 ± 5.67 ha) compared to the responsive cutters (34.6 ± 2.52 ha) or non-cutters (29.3 ± 5.35 ha) (Table 2). We found that the age of respondents in the non-cutter AFT were greatest and in the responsive cutter AFT were lowest, as measured by the age of the oldest and youngest of joint family owners. This was also reflected in ownership tenure (as of 2017), with the non-cutters having the longest (27.4 ± 1.70 years) and responsive cutters having the shortest (21.9 ± 0.83 years) tenures.

**Table 2:**
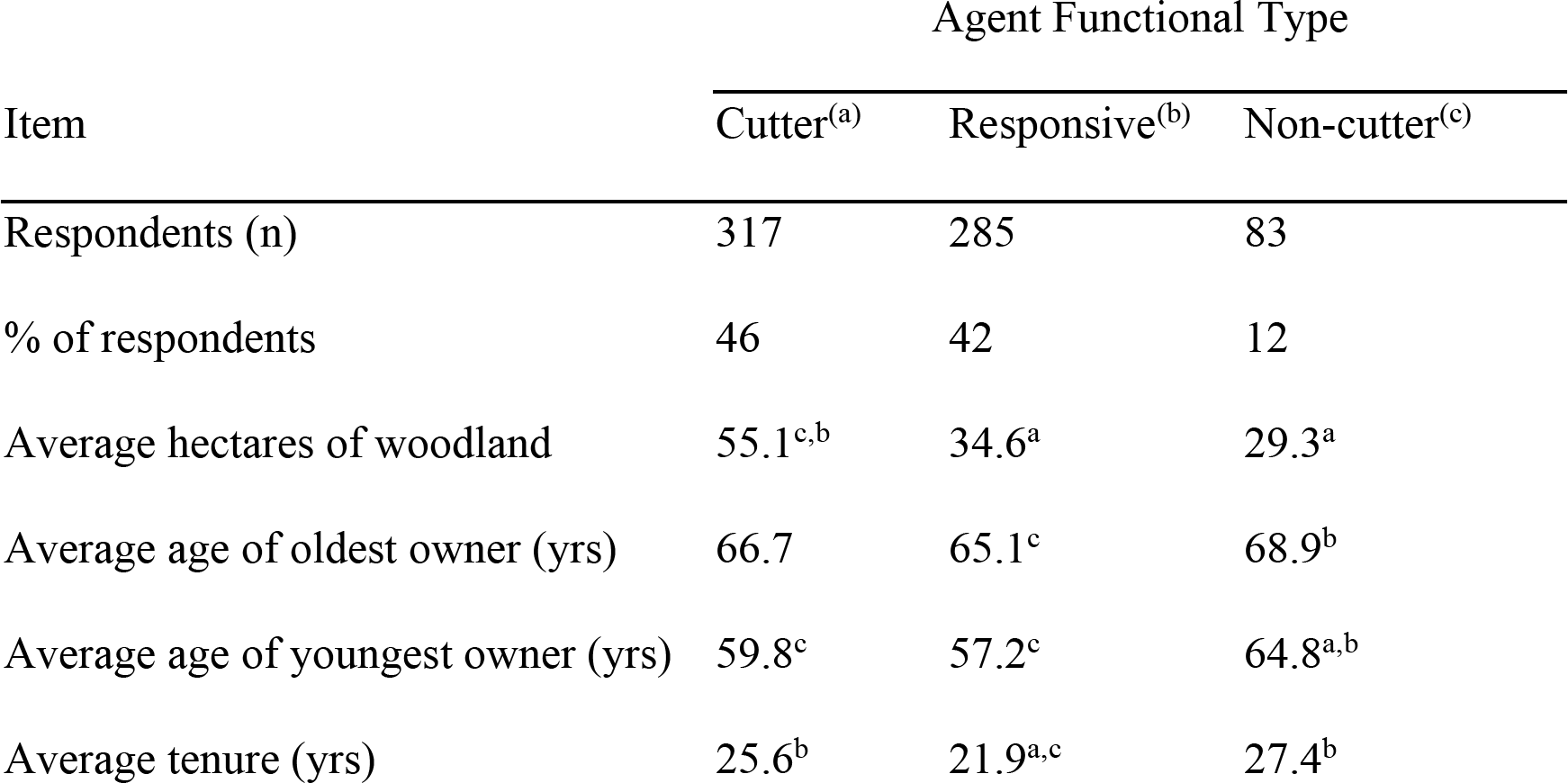
Number and percentage of FFOs, average hectares of woodland, average age, average tenure, and average number of owners by AFT. Superscript denotes statistical significance between (a) cutters, (b) responsive cutters, and (c) non-cutters in ANOVA posthoc Tukey significance test at the α = 0.05 level.

| Item | Agent Functional Type |  |  |
| --- | --- | --- | --- |
|  | Cutter<sup>(a)</sup> | Responsive<sup>(b)</sup> | Non-cutter<sup>(c)</sup> |
| Respondents (n) | 317 | 285 | 83 |
| % of respondents | 46 | 42 | 12 |
| Average hectares of woodland | 55.1<sup>c,b</sup> | 34.6<sup>a</sup> | 29.3<sup>a</sup> |
| Average age of oldest owner (yrs) | 66.7 | 65.1<sup>c</sup> | 68.9<sup>b</sup> |
| Average age of youngest owner (yrs) | 59.8<sup>c</sup> | 57.2<sup>c</sup> | 64.8<sup>a,b</sup> |
| Average tenure (yrs) | 25.6<sup>b</sup> | 21.9<sup>a,c</sup> | 27.4<sup>b</sup> |

**Table 3:** Characteristics of AFTs across gender, education, and household income.

| Attributes | Agent Functional Type (%) |  |  | Chi-square | P value |
| --- | --- | --- | --- | --- | --- |
|  | Cutter | Responsive | Non-cutter |  |  |
| Gender |  |  |  | 16.74 | < 0.001 |
| male | 0.72 | 0.74 | 0.51 |  |  |
| female | 0.28 | 0.25 | 0.49 |  |  |
| Education |  |  |  | 13.25 | 0.10 |
| < high school | 0.02 | 0.01 | 0.01 |  |  |
| high school | 0.33 | 0.22 | 0.32 |  |  |
| bachelor's degree | 0.26 | 0.29 | 0.27 |  |  |
| advanced degree | 0.29 | 0.35 | 0.33 |  |  |
| Household income |  |  |  | 15.37 | 0.05 |
| < \$25,000 | 0.10 | 0.05 | 0.03 | | |

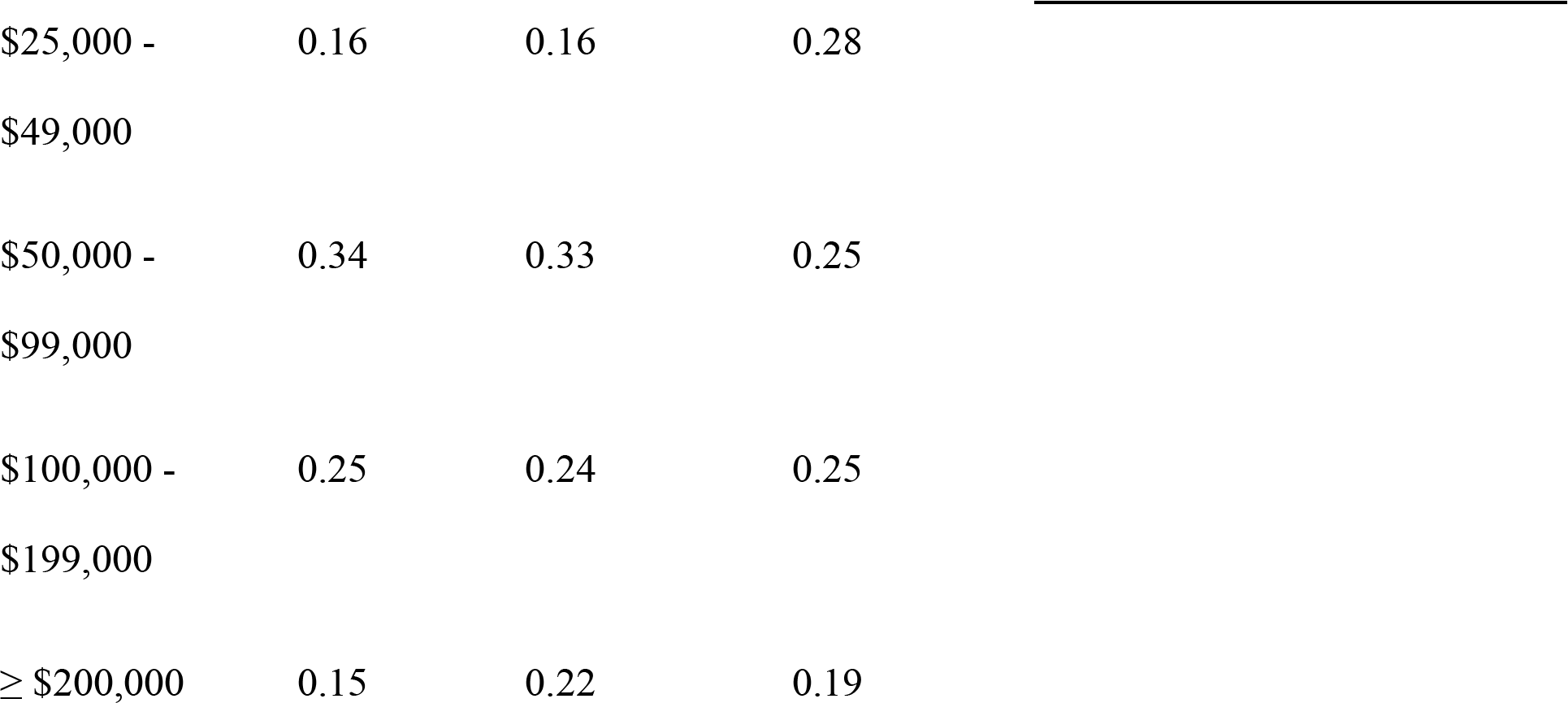

Cutters and responsive cutters were overwhelmingly male, whereas non-cutters were almost evenly split on gender (49% female compared to 51% male) (Table 3). Differences in educational achievement were greatest between cutters and responsive cutters; the cutter group had larger percentages of high school or lower education, and smaller percentages of bachelor’s degrees or advanced degrees, compared to the responsive group. The educational achievement of the non-cutter group fell in the middle of the cutters and responsive cutters. Annual earnings had a similar pattern between the cutter and responsive groups; the cutters had a greater proportion of low earners (< $25,000) and a smaller proportion of high earners (≥ $200,000) compared to the responsive cutters. Meanwhile, the non-cutters had the smallest percentage of low earners (< $25,000) but the highest percentage of earners in the $25,000 - $49,000 range.

Cutters, responsive cutters, and non-cutters differed in management history, building confidence in our functional typology (Figure 3). Cutters had a higher frequency of indicating that they had cut trees previously, sought advice from a forester, and had a management plan than the responsive or non-cutters. Furthermore, the responsive group fell between the cutters and non-cutters in all three questions.

**Figure 3:**
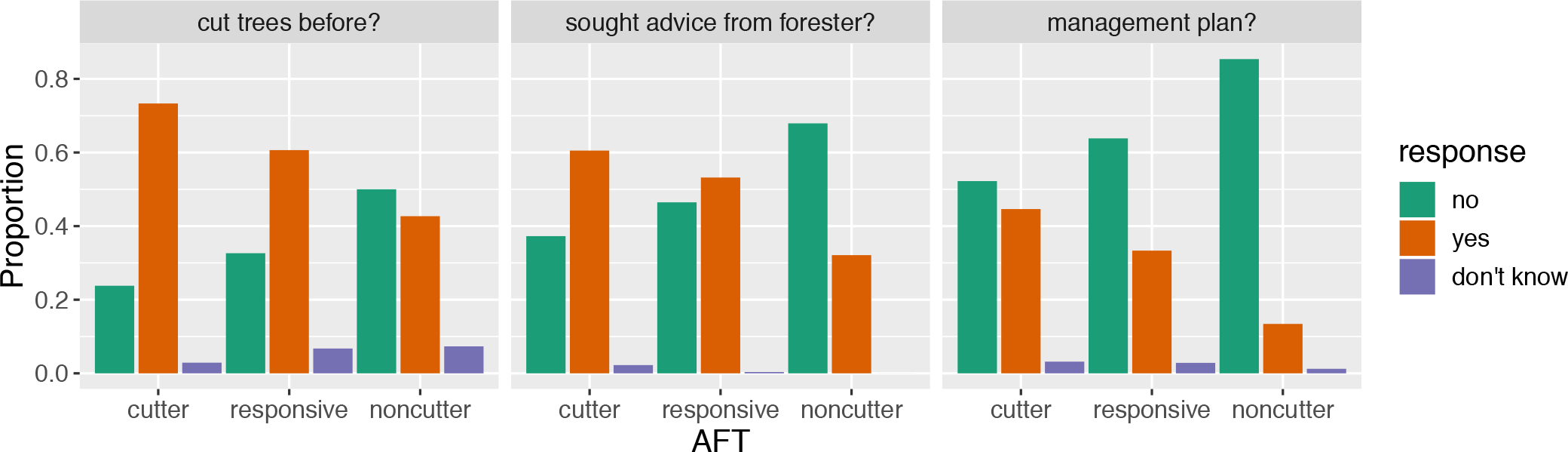
Proportion of each AFT with previous forest management experience.

The diversity of stated landowner objectives for ownership were simplified using PCA (Table 4), wherein the first four principal components captured approximately 70% of the variance. Component 1 weighs heavily on beauty, nature, and wilderness, which we label as "scenery". Component 2 is dominated by firewood, timber products, forest products, and hunting, which we call "utility". Components 1 and 2 reflect the cluster analysis of Majumdar et al. (2008), whose first two clusters were (1) “multiple-objective” and (2) “timber”. Components 3 and 4 are best represented by “investment” and “privacy”, respectively.

AFTs exhibit statistically significant differences across principal components 1 (scenery), 2 (utility) and 4 (privacy) (p < 0.001) (Figure 4). For both the scenery and utility scores, cutters had the highest values, followed by responsive cutters, then non-cutters. The privacy scores, on the other hand, were highest for the non-cutters and lowest for the cutters.

**Table 4:** PCA loadings of landowner objectives. Rotated component scores of the first four principal components > 0.30 are in bold.

| Landowner objective | Rotated principal component scores |  |  |  |
| --- | --- | --- | --- | --- |
|  | PC1: | PC2: | PC3: | PC4: |
|  | “scenery” | “utility” | “investment” | “privacy” |
| beauty | **0.43** | -0.27 | 0.05 | 0.07 |
| nature | **0.44** | -0.29 | -0.01 | **-0.37** |
| wilderness | **0.45** | -0.22 | 0.01 | **-0.39** |
| investment | 0.03 | 0.19 | **0.88** | -0.10 |
| privacy | 0.29 | -0.15 | **0.32** | **0.64** |
| heirs | 0.21 | 0.18 | -0.08 | 0.28 |
| firewood | 0.24 | **0.42** | -0.02 | 0.09 |
| timber | 0.15 | **0.47** | 0.14 | **-0.38** |
| products | 0.25 | **0.38** | -0.19 | -0.01 |
| hunting | 0.19 | **0.39** | -0.20 | 0.07 |
| recreation | **0.35** | 0.01 | -0.15 | 0.24 |

**Figure 4:**
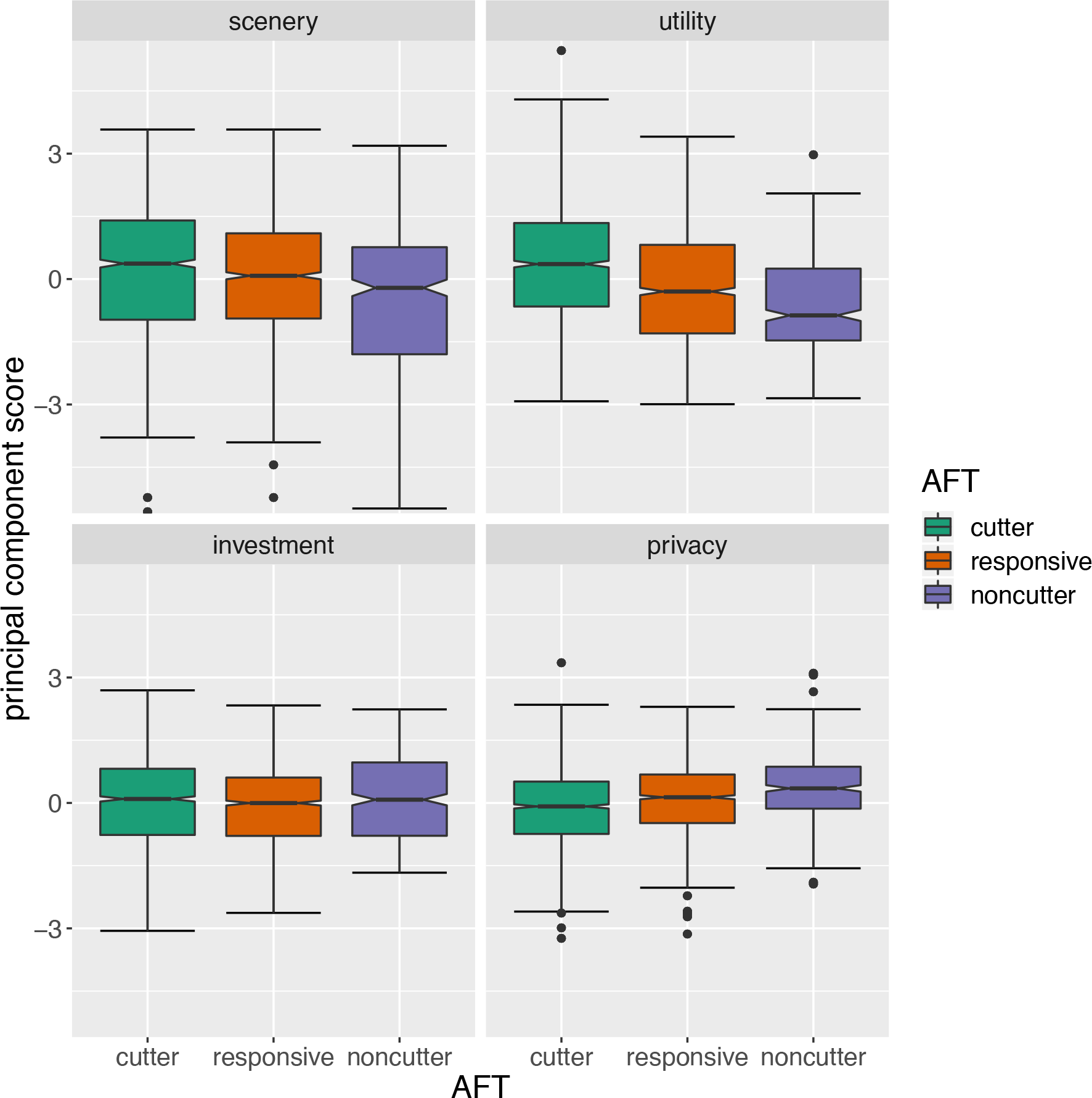
Boxplot of landowner objective principal component scores by AFT. One or more AFTs are significantly different from the others across principal components 1 (scenery), 2 (utility) and 4 (privacy).

The many available town-level socio-demographic variables were also simplified using PCA. The first two principal components of the town-level socio-economic variables explain approximately 60% of the variance and can be identified as (1) “wealth” and (2) “age” related dimensions (Table 5). AFTs exhibit statistically significant differences across both (p < 0.001) (Figure 5). Both cutters and responsive cutters score lower than non-cutters on the wealth principal component, indicating that landowners inclined to harvest trees live in towns that are poorer on average (higher fraction of town in poverty, lower median household income). In addition, cutters and responsive cutters score higher than non-cutters on the age principal component (higher median age, and higher social security and retirement income). It should be noted that FFOs are older on average than the general population (B. J. Butler et al., 2016), however the findings in Figure 5 provide an interesting juxtaposition to those in Table 2; while non-cutters are older on average than the other two groups, non-cutters tend to live in towns that are on average younger than the other two groups.

**Table 5:**
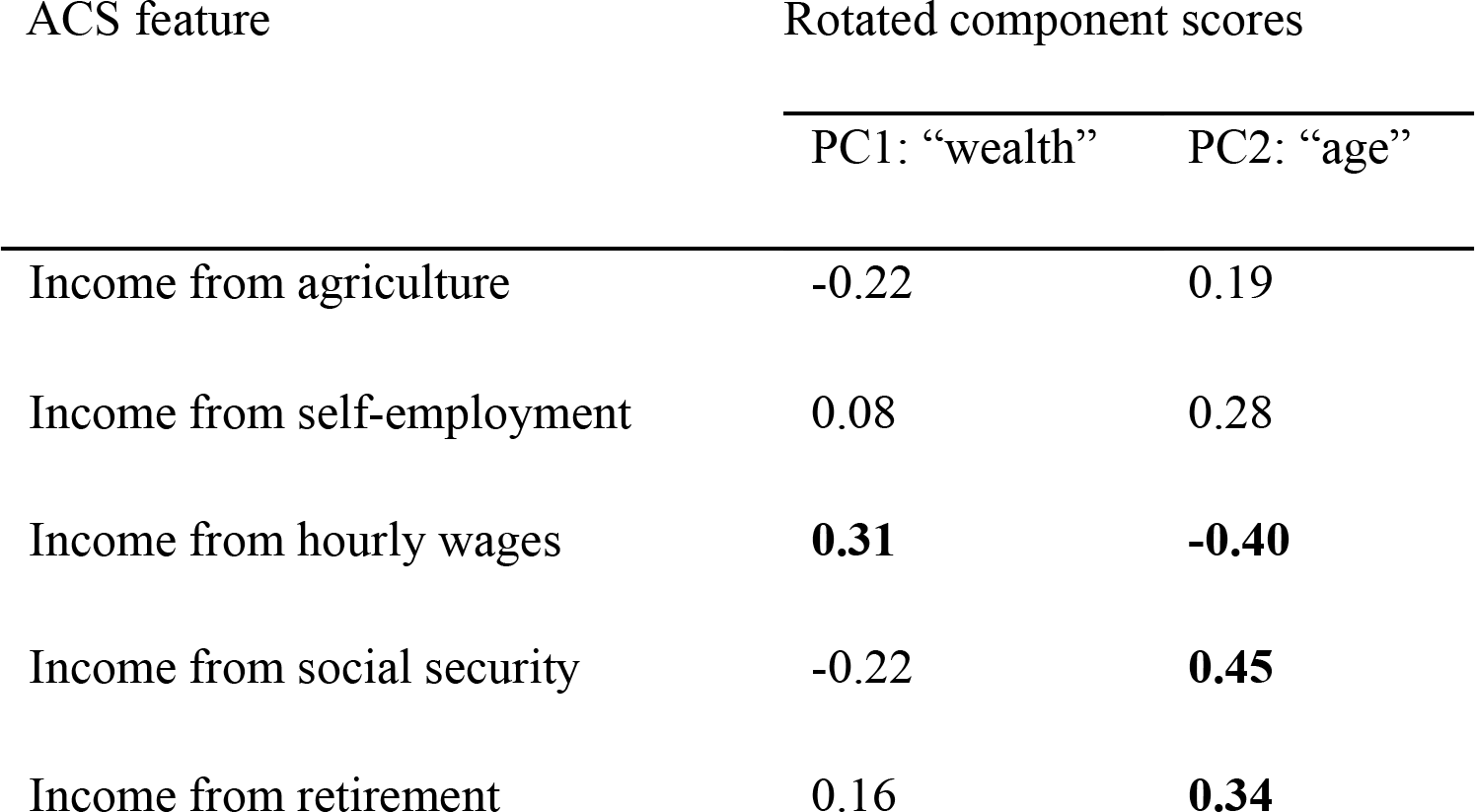
PCA scores of town socio-demographics. Rotated principal component scores > 0.30 are in bold.

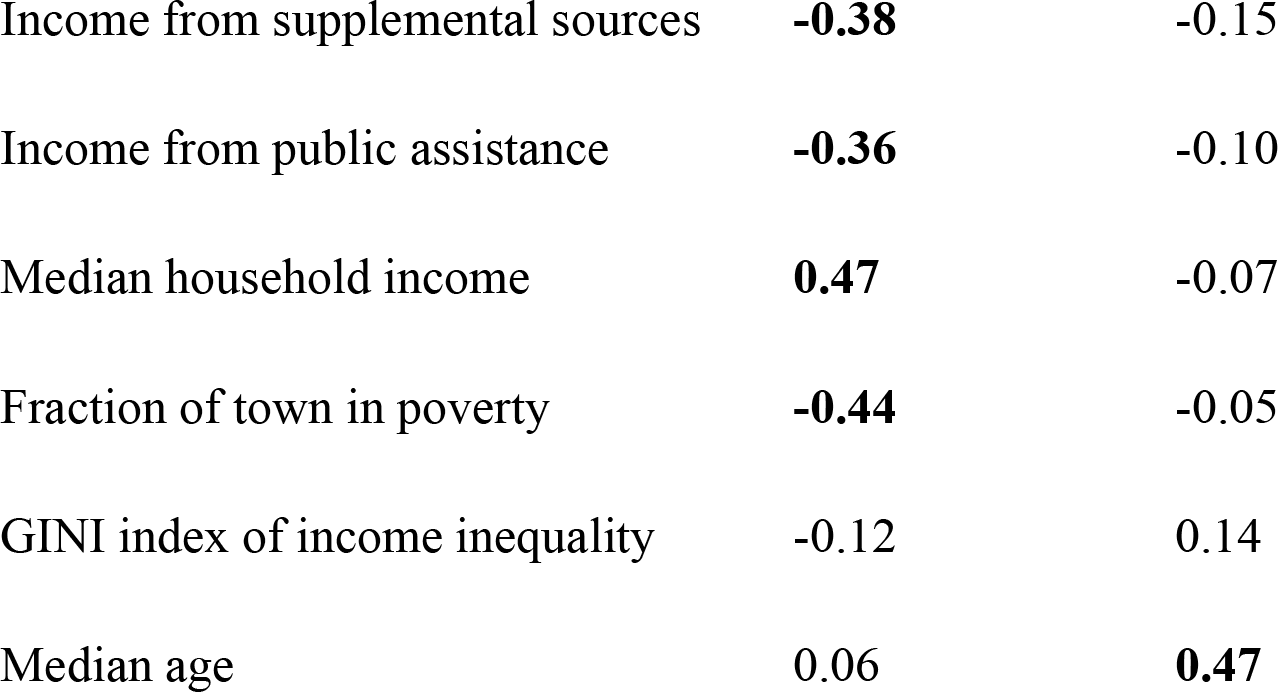

**Figure 5:**
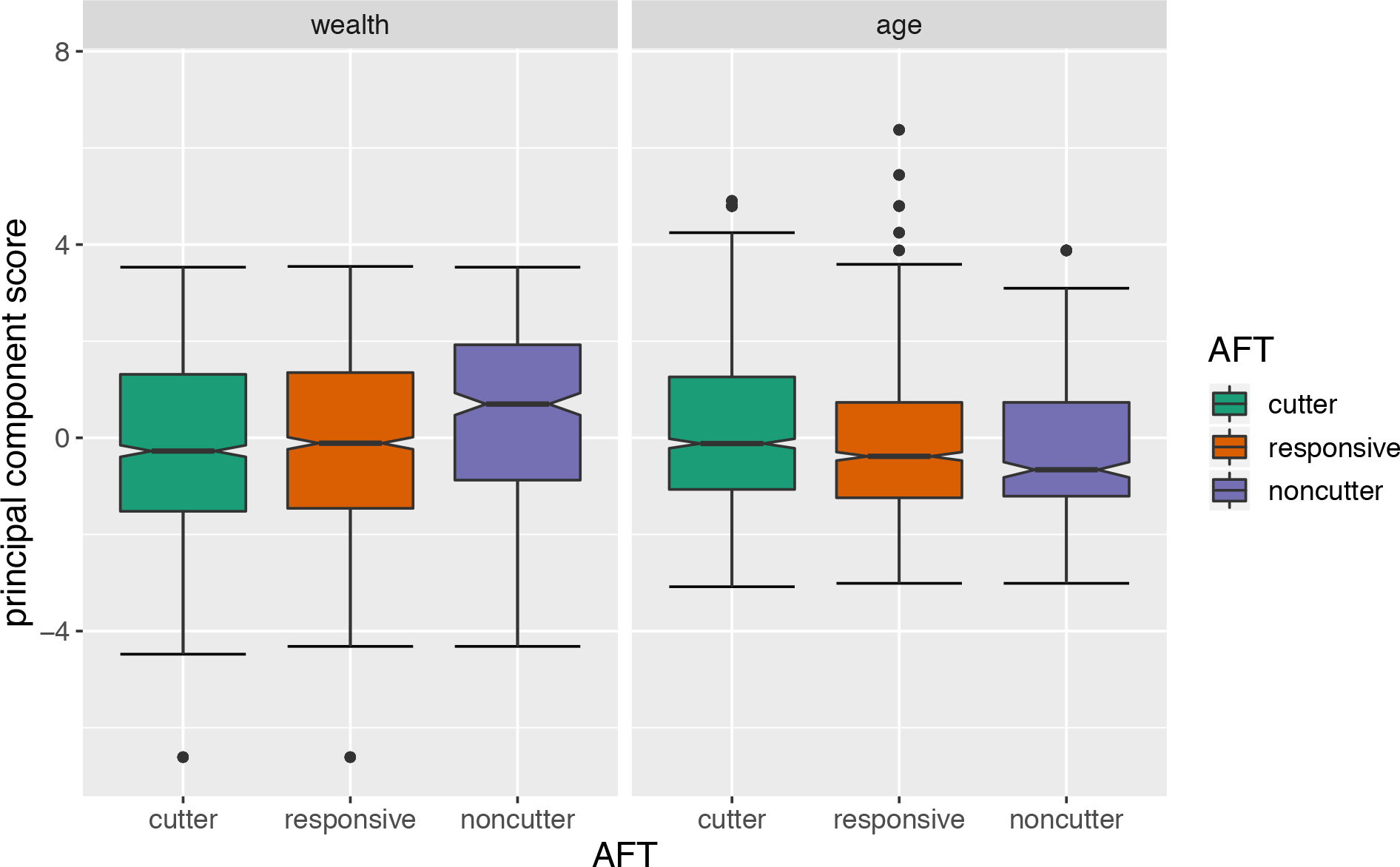
Boxplot of socio-demographic principal component scores by AFT. One or more AFTs are significantly different from the others across principal components 1 (wealth) and 2 (age) (p < 0.001).

### 3.3 Predicting AFTs Across the Landscape

A CART model fitting AFT to the survey stratification factors (Supplemental Information B) reveals a non-monotonic spatial trend in the distribution of AFTs. Small parcels (< 20 ha) in all regions except those in New Hampshire exhibit a significantly different distribution of AFTs than large parcels (≥ 20 ha) or those located in regions in New Hampshire (Figure 6). To incorporate this pattern into our modeling scheme, we constructed a new two-level factor variable called “Zone” from the 12 original survey strata: (A) parcels ≥ 20 ha plus all parcels in New Hampshire (north and south); (B) parcels < 20 ha in Vermont north and south, Massachusetts, and Connecticut.

**Figure 6:**
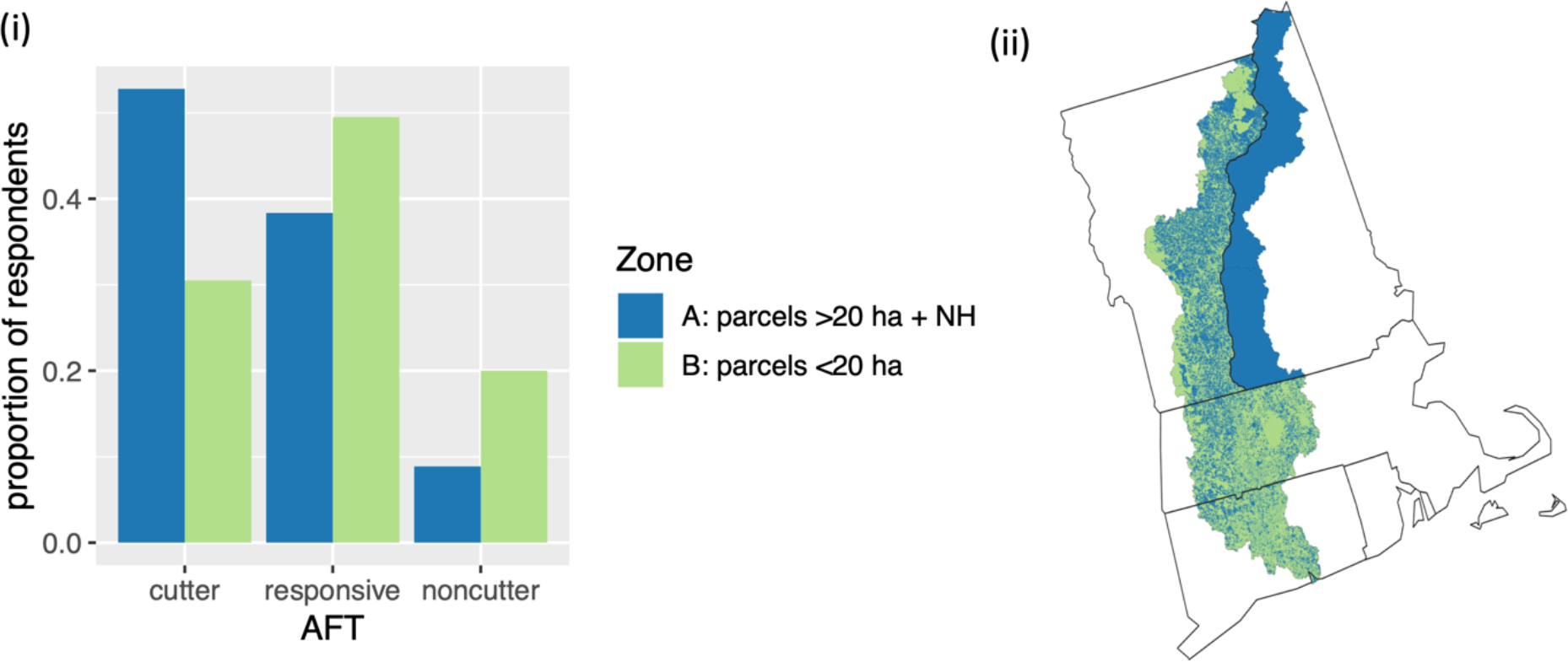
Distribution of AFTs across the study area. (i): “Zone”, a constructed two-level factor variable. Parcels ≥ 20 ha and all parcels in New Hampshire (blue) are predominantly cutters, whereas the remaining strata are predominantly responsive cutters (green); (ii): Zone A (blue) and Zone B (green).

We next fit a multinomial logistic regression model to predict AFT from “Zone” as well as from parcel-and town-scale geographic predictors. Stepwise AIC model selection reveals that the best model includes the following three predictors: Zone, town-level forested fraction, and parcel-level area of woodland (Figure 6). We also included an interaction term between Zone and the two continuous features (town-level forested fraction and area of woodland). The interaction term revealed that town-level forested fraction is significant with respect to Zone B (parcels < 20 ha), but not with Zone A (larger parcels + NH).

The model suggests that with increasing area of woodland owned, a respondent has an increasing probability of being a cutter or responsive cutter, and lower probability of being a non-cutter (Figure 7). As forest fraction increases in a town, we anticipate an increase in the probability of cutters and a decrease in the probability of responsive and non-cutters. The McFadden’s pseudo R^2^ of this model is 0.29. Model coefficients and standard errors are reported in Supplemental Information C.

**Figure 7:**
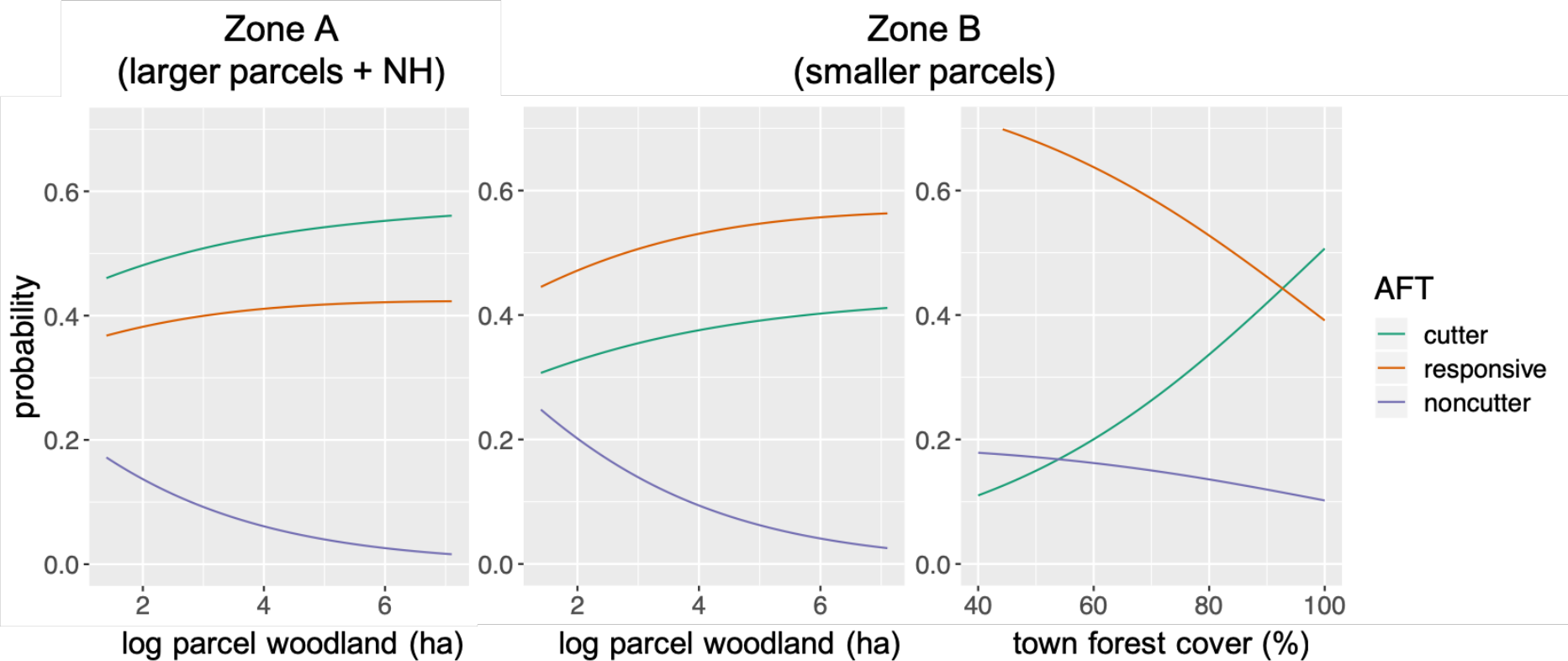
Summary of multinomial logistic regression results, with AFT probability expressed as a function of model predictors for each of the two zones. In each panel, variables not indicated on the horizontal axis were held at their mean value.

Using the multinomial logistic regression (Figure 7), AFTs were predicted for all parcels in the Connecticut River Watershed, including those not owned by survey respondents (Figure 8). The cutter and non-cutter probabilities exhibit distinct north-south trends, with notably more cutters in the north, whereas the responsive cutters are relatively evenly distributed throughout the study region. The responsive cutter group also has the highest probability overall.

**Figure 8:**
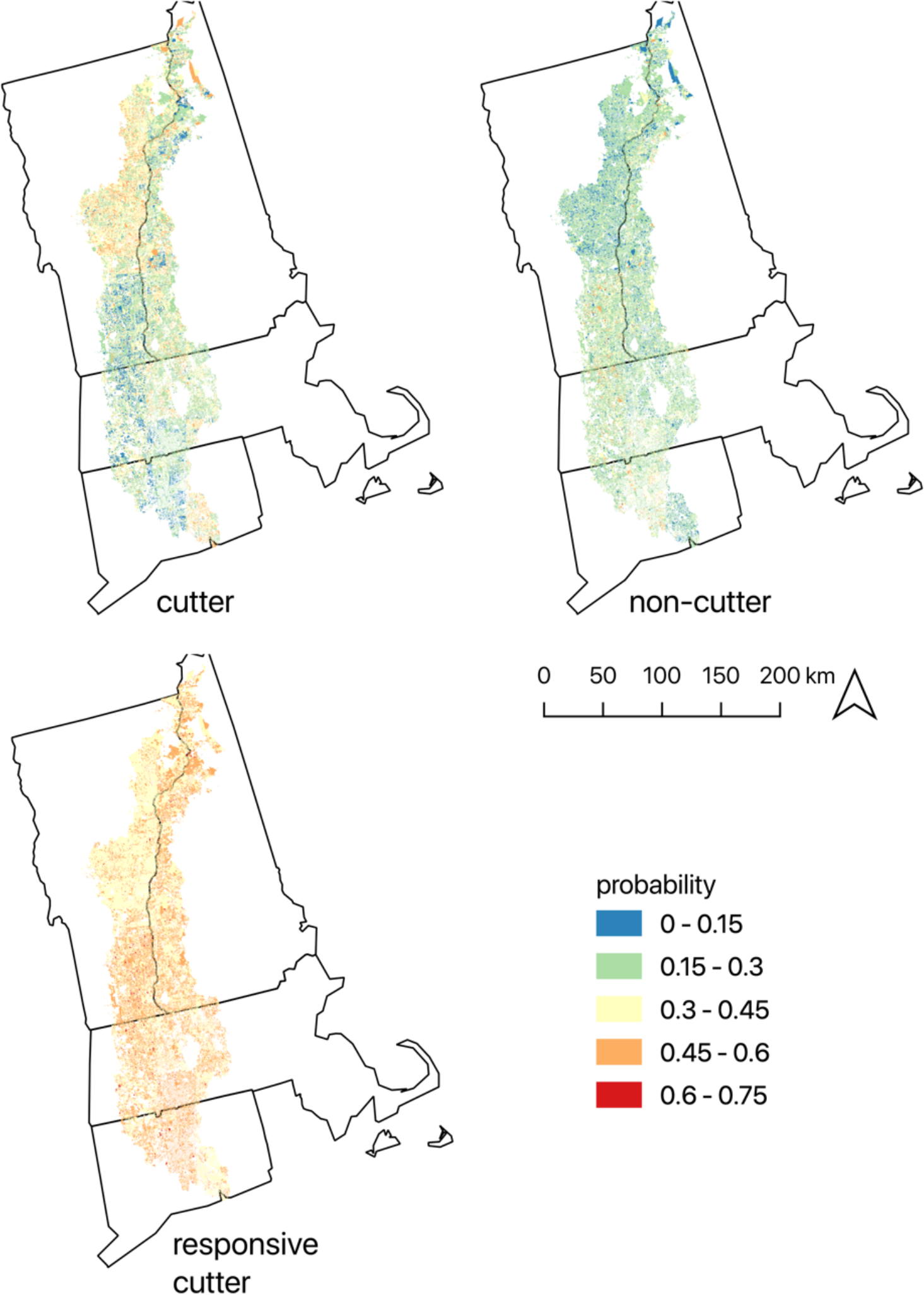
Predicted AFT probability throughout the Connecticut River Watershed, calculated from a multinomial logistic regression model based on zone, parcel-level hectares of woodland, and town-level forested fraction.

## 4. Discussion

As FIPs continue to spread in New England and disturb the forested landscape, an important question emerges: how will human behavior in response to FIPs interact with the disturbance created by these FIPs? And how will these interactions in turn impact the spread and impacts of FIPs? In our survey, 88% of respondents indicated that they would consider harvesting in response to FIPs, depending on the severity of the infestation. Just over half of the landowners reported having harvested their land before, indicating that FIPs may incite harvest on more parcels than would normally be harvested during routine forest management in the Connecticut River Watershed. Indeed, with our new knowledge regarding the behaviors of these different groups of landowners and our ability to predict AFTs and AFT behavior across the landscape, we may now be able to more holistically look at how FIPs disturb forests, inclusive of the management response to FIPs. Coupling human and FIP behavior is essential for modeling the ecological and economic impacts of invasive insects, particularly in New England, where there are > 200,000 forest landowners and among the highest numbers of FIPs in the U.S.

Clustering landowners based on behavior is useful because the AFTs translate naturally to models of landscape change in which human decisions are simulated alongside ecological processes (Arneth et al., 2014). While other studies have clustered private landowners based on survey data, our approach is novel for its use of empirical information collected through CBQs. Our clusters are *agent functional types* because they represent patterns of contingent *behavior*. Because we clustered based on contingent behavior alone (omitting objectives, management style, and demographics), our typology does not fall along clean lines of landowner objectives and motivations, suggesting that (a) tree cutting can be affected by a variety of motivations and (b) clustering on the basis of cutting behavior alone does not capture the range of landowner intentions. However, we chose to prioritize contingent behavior over descriptive characteristics because, in our case, understanding the behavioral response of landowners to FIPs is our end-goal. This AFT modeling framework can be directly applied to compute parcel-level probability of harvest under different FIP scenarios.

### 4.1 Cutters

In the survey, 73% of our cutter AFT indicated that they had previously cut trees on their property, and 45% indicated that they had a management plan - the highest in both categories of any AFT− as we would expect. The cutters own the largest fraction of the watershed (59.9%), and also own more woodland on average, suggesting that larger parcels provide more opportunity for sustaining economically viable timber harvests. Furthermore, this group exhibits the highest score on the “utility” principal component (firewood, timber, non-timber products, and hunting) as well as the “scenery” principal component (beauty, nature, wilderness, and recreation), indicating that landowner motivations to cut span multiple landowner objectives. Town-level ACS data suggest that while the cutters are not the oldest individuals, they are more likely to live in towns with older average populations, and also in towns with lower median incomes. This trend could be due to the fact that large swaths of woodland are located in older, lower wage-earning, more remote regions of New England. The cutters themselves are also more likely to earn < $25,000 and less likely to earn > $200,000 compared to the other groups. Cutters have the greatest potential to intensify the impact of FIPs on the landscape, as 46% of the respondents were grouped into this AFT, are always predicted to cut in response to a FIP, and generally own the largest parcels.

### 4.2 Responsive Cutters

The responsive cutter AFT was sensitive to variation in the parameters of the CBQs. Responsive cutter coefficient estimates on mortality percent and mortality years follow intuition: The greater the tree mortality percent (i.e., severity) resulting from the FIP infestation, the more likely these FFOs are to cut their trees. Additionally, the greater the time to tree mortality (i.e., delay of damage), the less likely these FFOs are to remove timber. It is unclear whether the responsive cutter decisions are driven primarily by financial (i.e., pre-emptive or salvage logging) or other (i.e., safety, aesthetic) motivations. However, survey results indicate that responsive cutters are less likely to have cut previously, are less likely to have a management plan, and have a higher likelihood of owning their land for privacy, compared to cutters. Furthermore, responsive cutters exhibit higher likelihoods of earning larger incomes and obtaining advanced college degrees. Therefore, we hypothesize that responsive cutters have different motivations for cutting compared to cutters, and this group’s lower ranking on the “utility” principal component perhaps suggests that responsive cutters are less financially-driven relative to cutters. Spatially, the responsive group is more likely on parcels < 20 ha and outside of New Hampshire, compared to the cutters and non-cutters. We frame the typical responsive cutter as a relatively young landowner with little past management experience who is willing to harvest infested trees, but unsure of the degree to which he or she will commit to doing so. Approximately 33% of the Connecticut River Watershed is controlled by the responsive cutters, making up 42% of the survey respondents. This group owns on average the second-largest parcels (after the cutters), and will, on occasion, harvest in response to FIPs, moderately increasing the impact of FIPs on the landscape.

### 4.3 Non-cutters

Non-cutters are our smallest group, encompassing only 12% of the survey respondents. They own the smallest area of forest, have the longest tenure, and are the oldest on average. Non-cutters are just as likely to be female as male, in stark contrast to the other two groups, which are predominantly male. This gender disparity reflects the global trend of lower rates of forest management among women, compared to male landowners (S. M. Butler, Huff, Snyder, Butler, & Tyrrell, 2017). The non-cutters are the least experienced in forest management activities, and also score the lowest on the utility objective principal component. Furthermore, the non-cutters score the highest on the privacy component, indicating that the landowners in this AFT value their land as a retreat rather than a source of revenue. While the non-cutters are the oldest individuals, they live in towns that are younger and wealthier on average, perhaps due to the fact they own smaller parcels, which can be found in less remote areas. This demographic – the non-cutters – are more inclined to let nature take its course through passive amenity appreciation and the least likely to amplify the impact of FIPs on forests.

### 4.4 Limitations and Future Research

Cutters and non-cutters by definition exhibit consistency in their responses to the CBQs, although it is important to consider these results only within the range of values presented to them. It is entirely possible that a cutter would choose not to cut under less severe circumstances than those presented in the survey, and that a non-cutter would respond differently under harsher conditions. Nevertheless, the range of values in the CBQ are reflective of realistic conditions.

Still missing from this puzzle of how FFOs will influence the total disturbance initiated by FIPs is the distribution and types of FIPs causing the response, types of harvests applied in response to FIPs, and forest types affected by both the FIP and harvests in response to FIPs. Our typology provides a basis for FFO response scenarios, although more research is needed to obtain finer insight into the predicted impacts of FIP-induced harvests. Of particular importance could be the phenomenon of “by-catch”, which refers to the non-host tree species being harvested along with the FIP-host species so that the harvest operation can be commercially viable. Another open question is the role of loggers in FIP-induced salvage logging; even if cutters and responsive cutters seek to salvage infected timber (and by-catch), would the marketplace me amenable to an influx of product? Furthermore, it is possible that state-imposed quarantines could further suppress the ability of loggers to turn a profit? Future modeling within this AFT framework will also depend on the evolution of AFTs over time. Changing landscape conditions (e.g., forest loss to development), ownership characteristics (e.g., shifts in parcel size), and change within and between FIP species will all contribute nuance to human-FIP interactions over time.

Understanding the direct and indirect impacts of FIPs requires a coupled natural-human systems-based perspective (Field, Dayer, & Elphick, 2017; Knight, Cowling, Difford, & Campbell, 2010) because, while FIP infestations induce human impacts on the natural system, the natural system, in turn, then evolves from the human-impacted natural system (Meurisse, Rassati, Hurley, Brockerhoff, & Haack, 2018). Our typology lends itself naturally to coupled human-tree-FIP modeling, which is the next stage of our team’s research. The goal is to support land management professionals in exploring alternative future scenarios with respect to forest stocks and FIP spread.

## Acknowledgements

This material is based on work supported by the National Science Foundation Coupled Natural and Human Systems Grant No. DEB-1617075. This research also partially supported by the Harvard Forest Long Term Ecological Research Program Grant No. NSF-DEB 12-37491.

