## Supplemental Information for "Landowner Functional Types to Characterize Response to Forest Insects"

**Supplementary Information**

A. Landowner objectives for ownership

Survey respondents were asked the following question:

*"How important are the following as reasons for why you* *currently* *own this woodland?"*

- To enjoy beauty or scenery
- To protect nature or biological diversity
- To protect or improve wildlife habitat
- For land investment
- For privacy
- To pass land on to my children or other heirs
- For firewood
- For timber products, such as logs or pulpwood
- For nontimber forest products, such as berries or maple syrup
- For hunting
- For recreation, other than hunting

Respondents indicated by checking boxes 1-5 where 1 = Not important; 2 = Of little importance; 3 = Moderately important; 4 = Important; 5 = Very important.

B. Classification tree of AFTs using survey strata. The numbers in the leaves represent the counts of cutters, non-cutters, and responsive cutters, respectively. Parcels ≥ 20 ha in all strata, and also parcels <20 ha in New Hampshire are more likely to be assigned "cutter" (left branch).

All other strata are more likely to be assigned “responsive” (right branch). The classification tree was used to construct a two-level categorical variable, “Zone”. The two levels of region are “Zone A” (left branch) and “Zone B” (right branch).

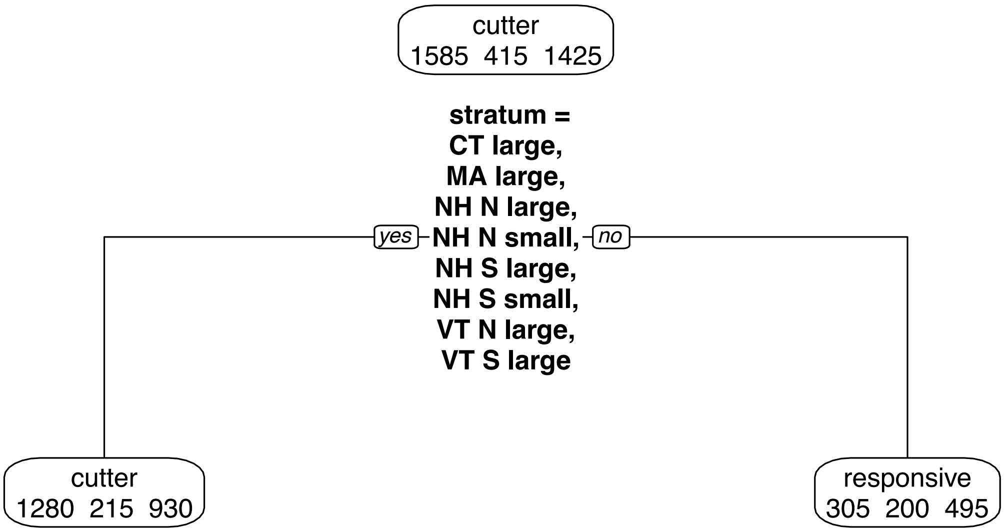


C. Multinomial logistic regression model coefficients and standard errors. Town forest cover is only in effect when Zone = B. Zone is defined in Supplemental Information B.

| AFT | Intercept | Region = smaller parcels | Log parcel woodland (ha) | Town forest cover (%) |
|---|---|---|---|---|
| coefficients | | | | |
| Cutter | 0.210 | -3.524 | 0.010 | 0.035 |
| Responsive | (reference level) | | | |
| Non-cutter | -0.145 | 0.126 | -0.441 | 0.001 |

| | | | | |
|---|---|---|---|---|
| Standard errors | | | | |
| Cutter | 0.029 | 0.116 | 0.008 | 0.001 |
| Responsive | (reference level) | | | |
| Non-cutter | 0.043 | 0.120 | 0.013 | 0.001 |
